# Functional analysis of flavivirus replicase by deep mutational scanning of dengue NS5

**DOI:** 10.1101/2023.03.07.531617

**Authors:** Amporn Suphatrakul, Pratsaneeyaporn Posiri, Nittaya Srisuk, Rapirat Nantachokchawapan, Suppachoke Onnome, Juthathip Mongkolsapaya, Bunpote Siridechadilok

## Abstract

Flavivirus NS5 is multi-functional viral protein that play critical roles in virus replication, evolution, and immune antagonism against the hosts. Its error-prone replicase activity copies viral RNA for progeny virus particles and shapes virus evolution. Its methyltransferase activity and STAT2-targeting activity compromise type-I interferon signalling, dampening protective immune response during infection. It interacts with several host factors to shape the host-cell environment for virus replication. Thus, NS5 represents a critical target for both vaccine and antiviral drug development. Here, we performed deep mutational scanning (DMS) on the NS5 of dengue virus serotype 2 in mammalian cells. In combination with available structural and biochemical data, the comprehensive single amino-acid mutational data corroborated key residues and interactions involved in enzymatic functions of the replicase and suggested potential plasticity in NS5 guanylyl transferase. Strikingly, we identified that a set of strictly conserved residues in the motifs lining the replicase active site could tolerate mutations, suggesting additional roles of the priming loop in viral RNA synthesis and possible strategies to modulate the error rate of viral replicase activity through active-site engineering. Our DMS dataset and NS5 libraries could provide a framework and a resource to investigate molecular, evolutionary, and immunological aspects of NS5 functions, with relevance to vaccine and antiviral drug development.

## Introduction

Flavivirus is a major group of arboviruses that cause a variety of human diseases transmitted by insects and mosquitoes. Several outbreaks by dengue viruses (DENV), Zika viruses (ZIKV), West-nile viruses (WNV), Japanese-encephalitis viruses (JEV), and yellow fever viruses (YFV) have caused significant morbidity and mortality worldwide (Pierson and Diamond, 2021). Dengue poses a major threat to public health in most tropical and subtropical countries. There are up to four-hundred-million infections per year and three-billion people are at risk of infection (Bhatt et al., 2013). Dengue vaccine development to account for all four serotypes of dengue viruses (DENV) has been a major challenge (Hadinegoro et al., 2015, Simmons, 2015, Torres-Flores et al. 2022).

Flavivirus NS5 functions as viral replicase and consists of three enzymatic activities located in two distinct domains. First, RNA-dependent RNA polymerase (RdRp) activity is responsible for viral RNA synthesis and resides in the C-terminal RdRp domain. Second, its methyltransferase (MTase) activities reside in the N-terminal MTase domain and methylate viral RNA at multiple positions such as the G cap at the 5’ end of viral RNA, the adenosine next to the cap, and the internal adenosines. N7- and 2’-O-methylation are two known forms of methylation imparted by flavivirus MTase. Third, the MTase domain contains guanylyl transfer (GTase) activity that adds GMP to the ppAG-RNA at the 5’ end in the first step of capping viral RNA with m^7^G cap. The m^7^G cap is required for binding with eIF4E during cap-dependent translation initiation. (Sahili et al. 2017)

In addition to copying viral RNA, flavivirus NS5 replicase can compromise both the early and the late phases of the type-I interferon (type I IFN) signaling. 2’-O-Methylation (2’- O-Me) of viral RNA shields it from the host surveillance system, preventing the launch of type I IFN response (Dong et al. 2012, Daffis et al. 2011, Züst et al. 2013). Flavivirus NS5 can bind and target human STAT2 (hSTAT2) for degradation, dampening the stimulation of hundreds of interferon-stimulated genes (ISGs) that confer antiviral state in cells (Best 2016). In addition to targeting STAT2, NS5 also interacts with multitude of host proteins to modulate cellular environment to support virus replication (Shah et al. 2018, De Maio et al. 2016, Bhatnagar et al. 2021).

Deep mutational scanning (DMS) is a massively parallel technique that could measure the functional effects of every possible amino-acid change at each position in a protein, with the ability to probe thousands of mutations (Fowler and Fields, 2014). The functional effects of mutations are measured in the form of a fitness score that reflects the frequency change of each mutant in a mutant pool (library) after a selection pressure is imposed on the library. Here, we applied DMS to probe the effects of all single amino-acid NS5 mutations on DENV viability in mammalian cells. We correlated our DMS data with structural and biochemical data of flavivirus NS5 to gain further insights into the enzymatic functions of flavivirus NS5.

## Results

### Deep mutational scanning in Vero cells

We used an infectious-clone plasmid system to construct the virus library of DENV2 strain 16681. The NS5 gene library was first constructed on pUC57 plasmid and then subcloned to substitute for the NS5 gene on the infectious-clone plasmid (Supplementary Figure 1). The infectious-clone plasmid contains an intron insert to stabilize the full-length flavivirus genome sequence in *E.coli* (Suphatrakul et al., manuscript in preparation). The comparison between the NS5 gene library and the infectious-clone plasmid library showed good transfer coverage of the NS5 variants from pUC57 plasmid system to the infectious-clone plasmid system, indicating that the potential genetic bottleneck from mutant instability in *E.coli* was minimal with the infectious-clone plasmid system (Supplementary Figure 2A). To accommodate the short-read deep sequencing, we broke down the NS5 library into 24 subpools with target mutagenic regions of 40 amino-acid length. The library was designed to contain 17,100 single amino-acid substitution mutants (saturation mutagenesis across all the 900 residues of NS5) and 1,333 synonymous mutants.

We probed the mutational effects on virus viability in a cellular setting with defective innate immune response to measure the “basal fitness” of the mutant viruses. An infectious-clone plasmid library of DENV2 mutants was transfected into BHK21-rtTA3 cells (a cell line with defective type-I IFN response) and the supernatant from the transfected cells was used to infect Vero cells. The mutant virus library from Vero cells was then harvested for amplicons preparation and deep sequencing (Figure 1A). We calculated weight-averaged fitness of mutants using DiMSum pipeline which estimated replicate-specific error to weigh the fitness scores across the replicates for averaging (Faure et al. 2020) (Supplementary Figure 3).

**Figure 1:**
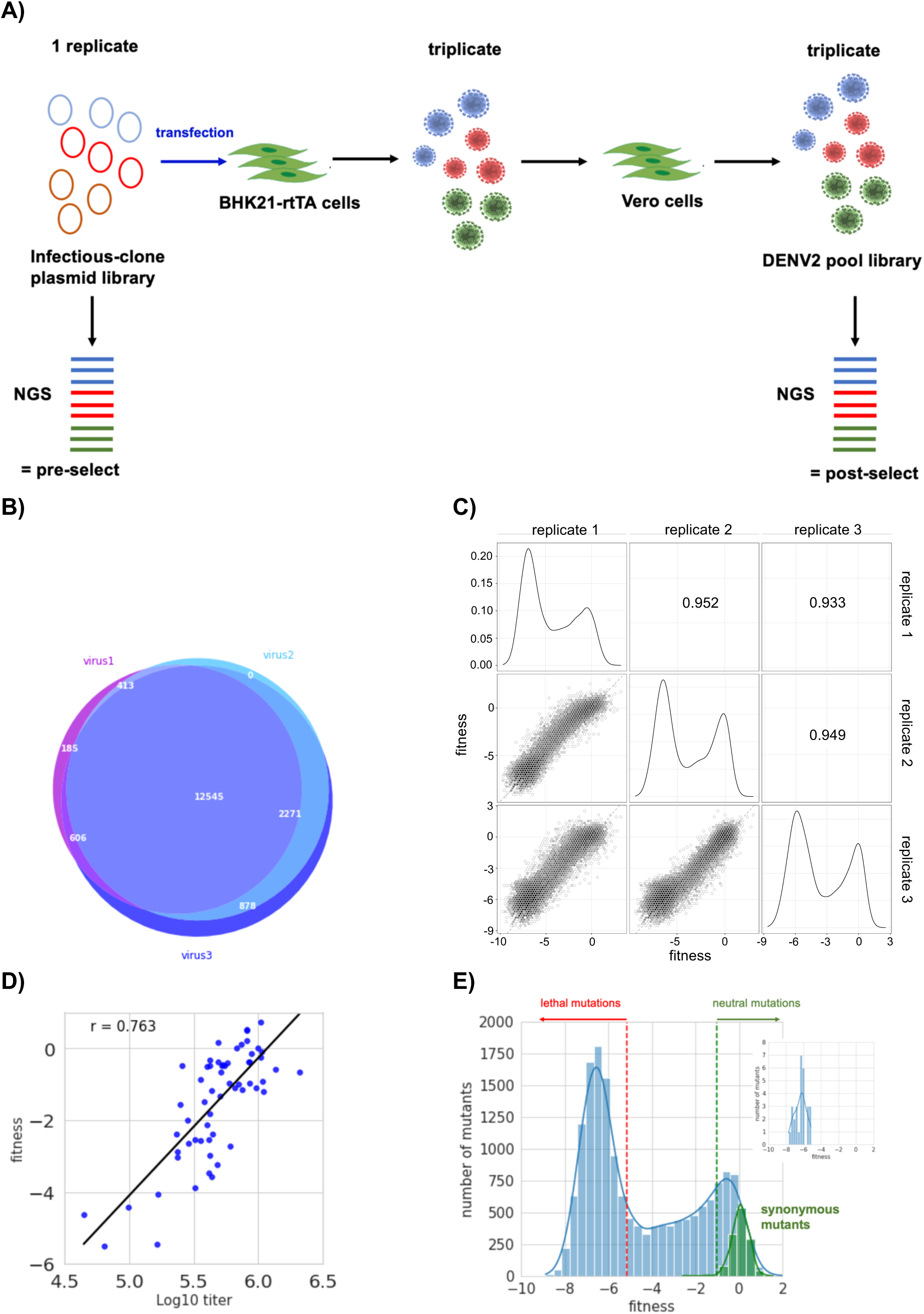
Deep mutational scannings of DENV2 library. A) Simplified diagram of DMS of DENV2-16681 NS5 library in Vero cells. B) Venn diagram showing the overlap of virus mutants with read counts more than zero among the three replicates of the DENV2 virus library obtained from Vero cells. The numbers represent the numbers of mutants in the corresponding areas on the Venn diagram. C) Correlation of DENV2 mutant fitness scores among the three replicates. D) Correlation between the fitness scores of selected DENV2 mutants and their corresponding infectious titers (2 days post infection) from A549 cells. E) Histogram of weighted average fitness scores of DENV2 mutants (blue = single amino-acid mutants, green = synonymous mutants). The green vertical dashed line defines the threshold of neutral mutants, while the red dashed line defines the fitness threshold of lethal mutants. The inset histogram shows distribution of fitness scores of 29 published lethal mutants used to define lethal mutation threshold (Supplementary table 1).

Deep sequencing of the infectious-clone plasmid library (input) showed mutant coverage with normal distribution (Supplementary Figure 1B). 99% of the designed single amino-acid mutants passed the input read-count filter during analysis (203 mutants were excluded from calculation of average fitness). Deep sequencing of the three DENV2 libraries (output) from Vero cells showed 80% overlap of mutants with read counts of at least 1 among the three replicates while over 80% of nonoverlapping mutants have read counts less than 4 in all the replicates, indicating no significant coverage bottleneck in our deep mutational scanning experiments (Figure 1B). The fitness scores of mutants between the three replicates were also reproducible, with high Pearson correlation of 0.95-0.93 among the three replicates (Figure 1C). Individual experimental measurements of virus titers at 2 days post infection in A549 cells infected with 63 mutant viruses showed strong agreement with their averaged fitness scores derived from the DMS dataset, with Pearson correlation of 0.76 (Figure 1D). In addition, our data also agree well with the previous mutational analyses of DENV2 (Supplementary table 1, and references therein). Based on 29 known lethal DENV2 mutations (Supplementary table 1 and references therein and Figure 1E), we identified 8791 mutations (or 52% of mutations covered by our DMS) with fitness lower than the maximum fitness score of the published DENV2 lethal mutations (Figure 1E). Based on the fitness score distribution of the synonymous mutations (Figure 1E), we identified 3005 neutral mutations (or 17.8% of mutations covered by our DMS). Thus, our DMS measurement showed high consistency among the three replicates and agreed with the experimental measurements.

### Mutation tolerance, sequence conservation, and amino-acid preference of dengue NS5

To assess mutational tolerance across NS5, we calculated mutation tolerance of each residue by averaging the fitness scores of all the single amino-acid mutations that pass the input read-count filter (Materials and methods). The heatmap plot (Figure 2A) and the mutation tolerance (top bars, Figure 2A) mapped onto a dengue NS5 structure (Figure 2B) show that the RdRp domain has strong mutation intolerance while MTase domain is relatively tolerant to mutations. Though the mutation tolerance is generally correlated with sequence conservation (represented by ConSurf score (Armon et al. 2001), high positive = high sequence variation, high negative = strictly conserved) of flavivirus NS5 (Pearson correlation = 0.72), there are multiple residues that deviate from the trend (Figures 2C). Calculation of the deviation between sequence conservation and mutation tolerance (i.e. difference mapping, Materials and methods) identified both the residues which did not tolerate mutation well despite low sequence conservation (high positive deviation, blue residues in Figure 2D) and the residues which tolerated mutations despite strict sequence conservation (high negative deviation, red residues in Figure 2D). We examined the nature of the amino-acid differences between the sequence conservation by multi-sequence alignment (MSA, Supplementary Figure 4) and the DMS data (Figure 2B) by calculating amino-acid preference of each residue (Bloom et al. 2014) (Figure 2A, WebLogo rows). We utilized the DMS data (Figures 2A-2B), amino-acid conservation (Figure 2B, Supplementary Figure 4), the deviation scores (Figure 2D), and amino-acid preference (Figure 2A) to examine functional sites of flavivirus NS5.

**Figure 2:**
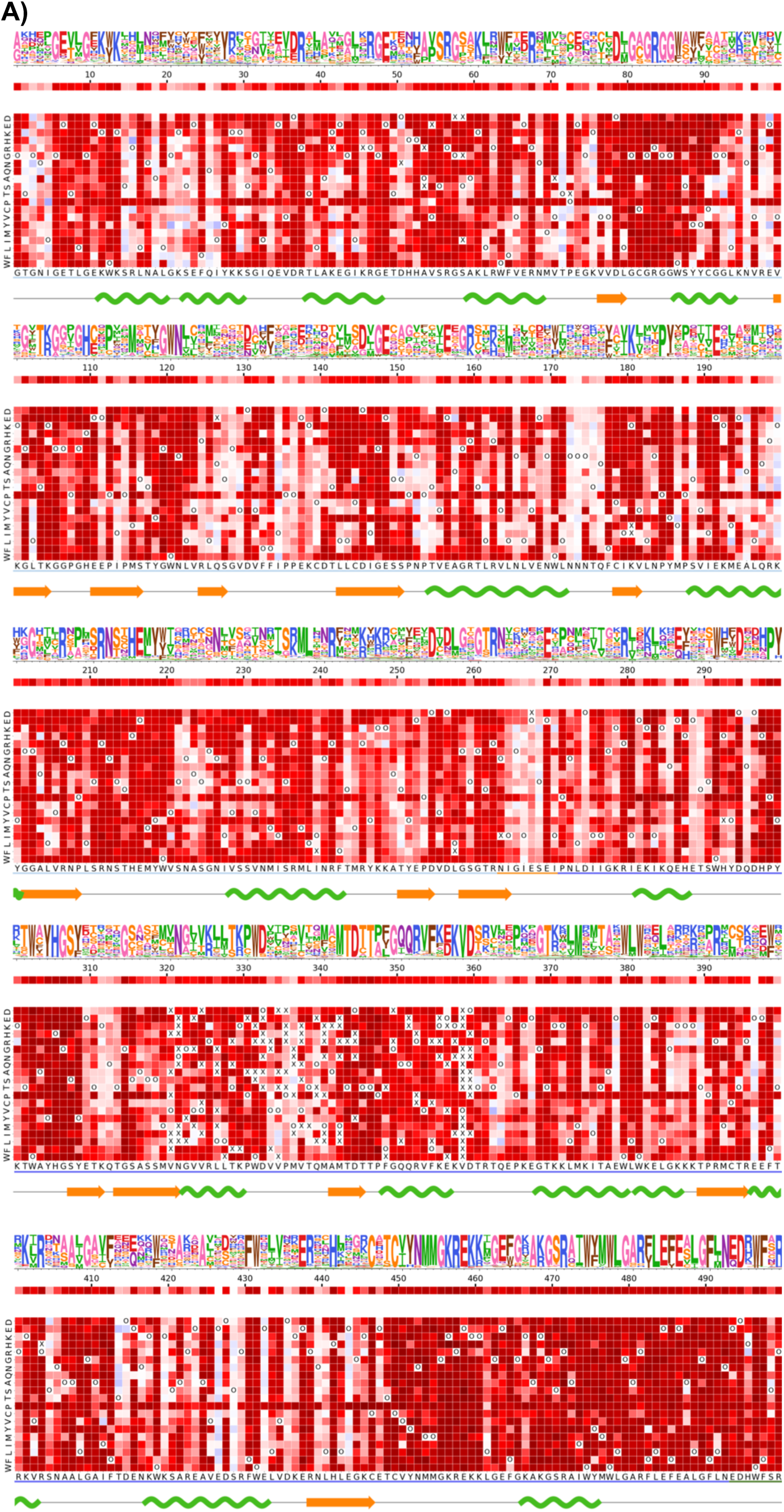
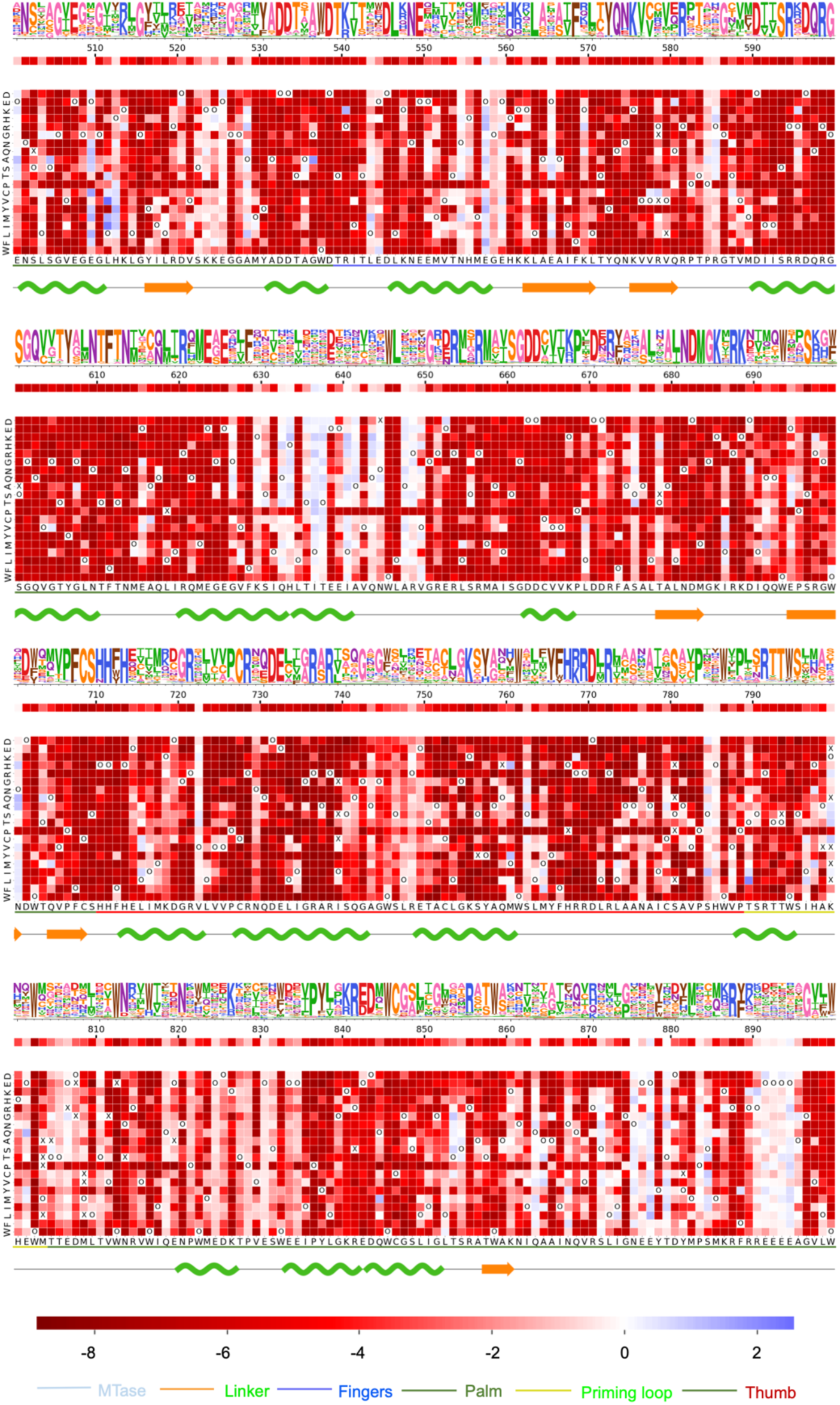
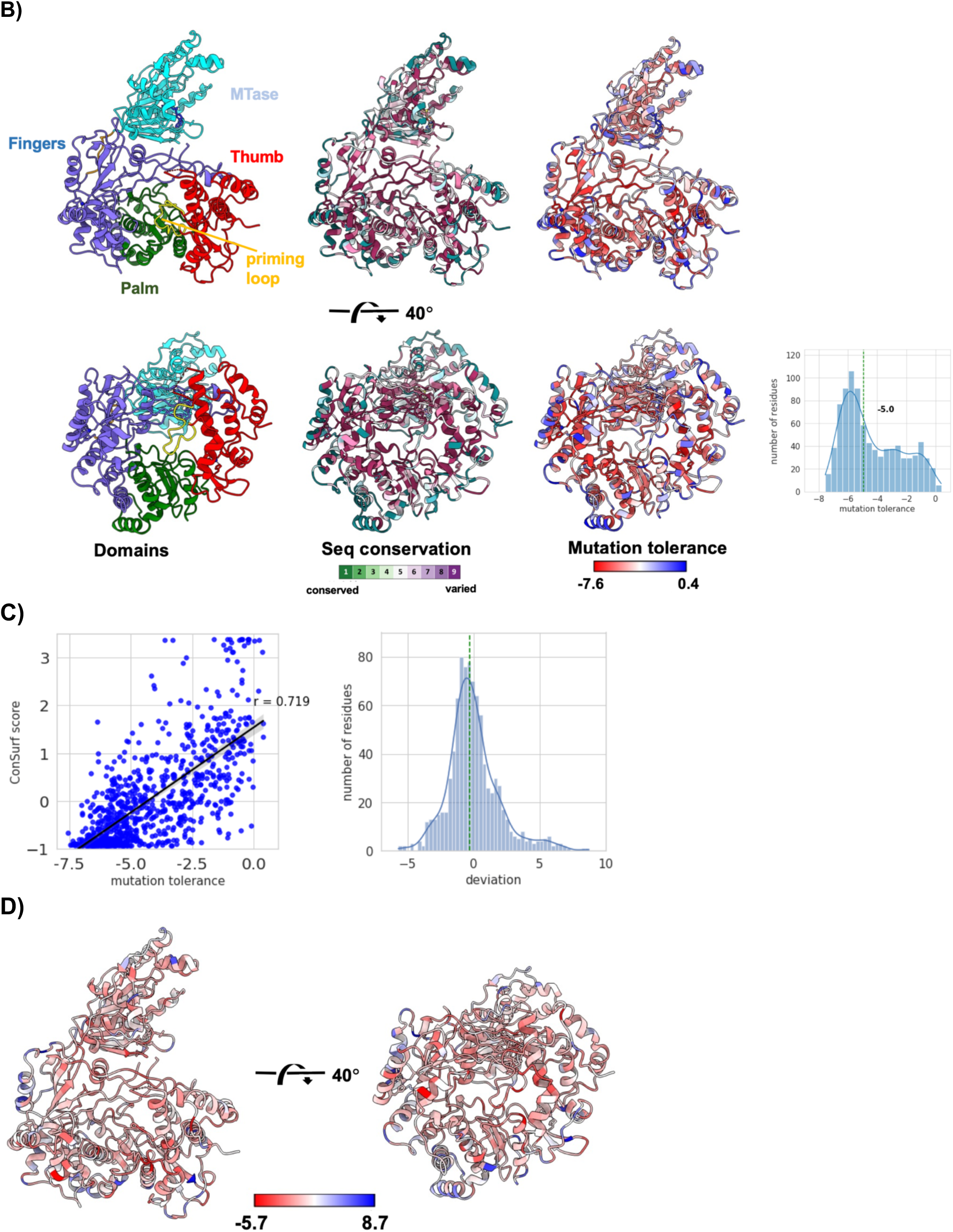
Fitness of DENV2 NS5 mutants, mutational tolerance, and amino-acid preference at each residue. A) WebLogo plot (top rows), heatmap of mutational tolerance (second row), and heatmap of mutants (third row) of each residue. DENV2 NS5 amino-acid sequence is labelled on the X-axis of the heatmap with the underline colored according to the domain that each residue is located. The “O” in the bottom heatmap marks the wild-type amino acid, while “X” marks the mutants that were filtered out by input read counts. Secondary structure of the NS5 is delineated on the bottom row where alpha helix is represented as wiggly line and the beta sheet is represented as arrow. The color bar for heatmap scores is shown at the bottom of the figure along with the color code for NS5 domains/regions. B) Structural mapping of sequence conservation (ConSurf, Ashkenazy et al. 2016) and mutational tolerance onto NS5 structure. The inset histogram shows the distribution of mutational tolerance scores. C) Correlation between the sequence conservation (ConSurf scores) and the mutational tolerance. Pearson correlation coefficienct = 0.719. Histogram of the deviation scores (the difference between the ConSurf scores and the mutational tolerance) is shown on the right. D) Structural mapping of the deviation scores on the NS5 structure. The structure of NS5 is based on the DENV2 NS5 by Wu et al. 2020 (6kr2.pdb).

### Analysis of the active site of NS5 RdRp domain

Our DMS data are consistent with several known characteristics of flavivirus NS5 RdRp domains (Malet et al. 2007, Yap et al. 2007, Surana et al. 2014, Godoy et al. 2017, Zhao et al. 2017, Dubankova et al. 2019, Yang et al. 2021). Deviation scores between sequence conservation and mutation tolerance are relatively low in most parts of the motifs A-G that make up the RdRp active site (Figure 3A). The catalytic D533 in motif A and the GDD in the motif C have strict amino-acid preference and conservation (Figure 3B). The motif C is strictly conserved and has absolute amino-acid preference (Figures 3A-B).

**Figure 3:**
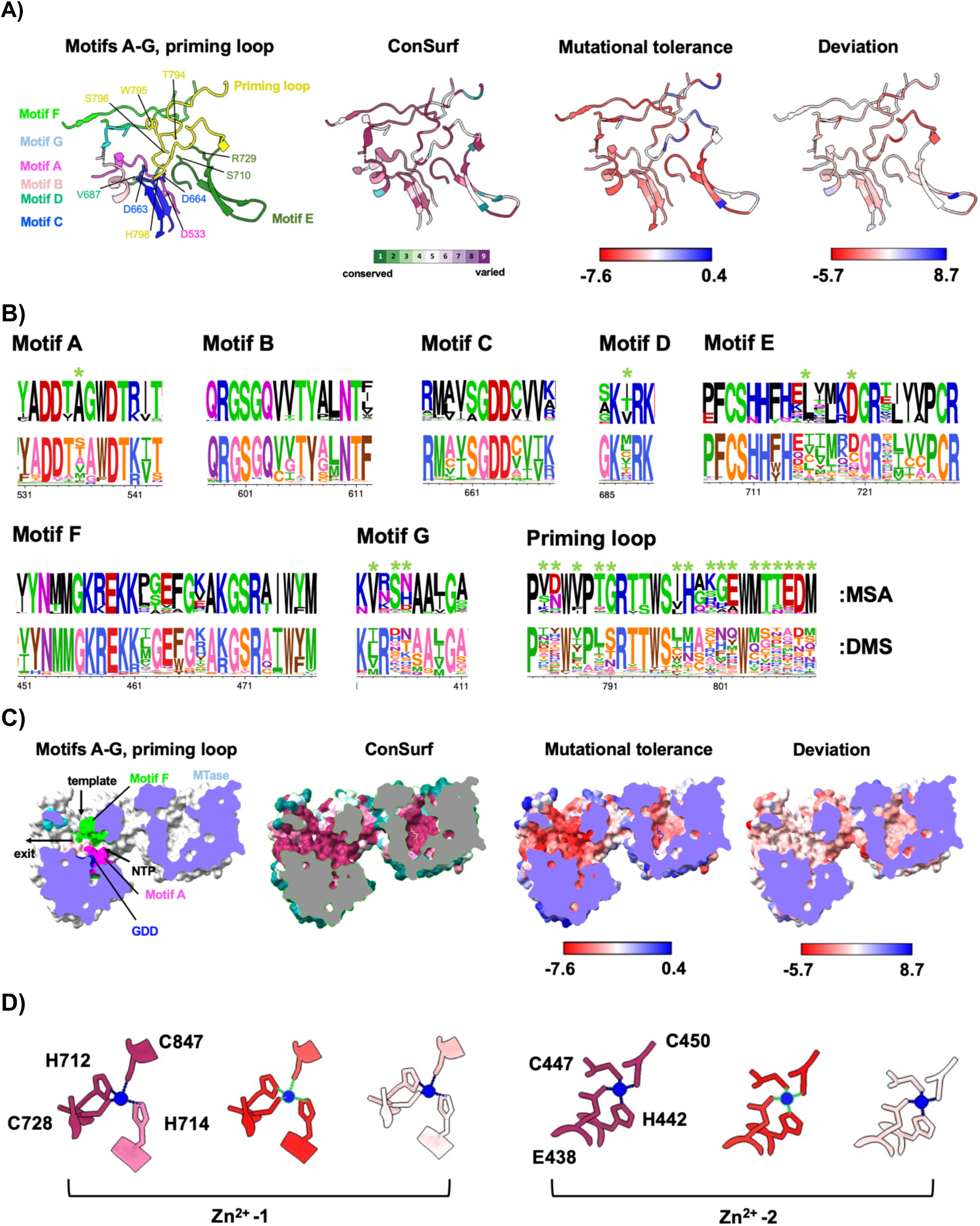
Analysis of mutational tolerance and sequence conservation of the RdRp active site. A) Structural mapping of ConSurf score, mutational tolerance, and deviation score on the Motifs A-G and the priming loop. B) WebLogo plots of multi- sequence alignment (MSA) of 92 representative flavivirus NS5 (top rows) and of the amino-acid preference calculated from DMS (bottom rows) of the Motifs A-G and the priming loop. The green asterisks mark the residues that tolerate mutations more than the sequence conservation suggests. C) A cross section of surface representation of the NS5 active site showing the template, the NTP, and the exit channels. From this view, the Thumb domain is facing the viewer and has been clipped out. D) Structural mapping of ConSurf score (left), mutational tolerance (middle), and deviation score (right) on the residues that coordinate with the two Zn atoms. The structure of NS5 is based on the 6kr2.pdb by Wu et al. 2020.

Three residues S710, R729, R737 that bind to *β*- and *γ*-phosphates of initiating ATP have strict amino-acid preferences and conservation (Figures 3A, 3B). The conserved priming-loop residue T794 and S796 that bind to the *α*-phosphate of the initiating ATP could accept certain amino acids but with much less preference to S and T (Figure 3B). Two conserved residues W795 and H798 were hypothesized to position the initiating ATP for *de novo* RNA synthesis. *In vitro* RNA assay showed that H798A reduced the formation of starting pppAG dinucleotide by half while W795A had the wild-type activity (Selisko et al 2012). In contrast, our DMS showed that W795 had strong preference for aromatic amino acids but H798 could be exchanged to several amino acids (albeit with lower preference for A) (Figure 3A). H798 has higher mutation tolerance than W795 (Figure 2A). This discrepancy suggests that W795 and H798 could be involved in additional steps of viral RNA synthesis. While most known mutations in the priming loop specifically affect *de novo* initiation but not elongation, E802A/Q803A (DENV3/DENV4) could also affect elongation activity (Selisko et al. 2012, Lim et al. 2016). The priming loop is hypothesized to retract from the active site after initiation to allow the occupation of both template and daughter RNA strands during elongation (Appleby et al. 2015). Thus, its dynamics may influence overall viral RNA synthesis.

Strikingly, despite strong sequence and structural conservation of motifs A-G and the priming loop that makes up the replicase active site among flavivirus (Peersen 2019), we identified multiple residues that could tolerate mutations in these motifs (Figures 3A- B). In particular, motif G, which has been implicated in regulating translocation during nucleotide addition cycle (Wang et al. 2020b), could accommodate several amino acids at most positions (Figure 3B). Motif D V687 could also tolerate multiple mutations (Figures 3A-B). Previous studies with poliovirus RdRp have implicated the Motif D in both catalytic and fidelity of nucleotide incorporation (Yang et al. 2012, Liu et al. 2013). In addition, the priming loop could also accommodate diverse mutations at several positions (Figures 3A-B). These highly mutable, yet-conserved residues in the active sites of flavivirus RdRp might play important roles in determining the fidelity of the viral replicase. Translocation speed of RNA polymerase could affect error rate of transcription (Gamba et al. 2018). While the reduction of error rate of a viral replicase could affect the virus natural fitness in nature and *in vivo* as the virus would be less capable of adaptation, it should be less critical for the viral fitness in a more homogeneous, experimental condition such as virus culture using Vero cells.

In addition to the active site, the template and the NTP channels have low mutational tolerance and high sequence conservation (Figure 3C). Two Zn^2+^ atoms have been consistently observed in the available flavivirus RdRp structures. The residues that coordinate with the first (H712, C728, C847, H714) and the second Zn^2+^ (C447, C450, H442, E438) have strict amino-acid preference, were among the residues with the highest mutation intolerance in NS5 and have strict amino-acid conservation (Figures 2A and 3D).

### Analysis of functional sites of NS5 RdRp domain

NS5 contains additional functional sites important for replicase function. Several studies have implicated the roles of interactions between NS5 and structural elements on viral RNA (vRNA) in flavivirus replication. In particular, surface accessible, positive-charge residues are candidates for the NS5-vRNA interaction. NS5 binds to the stem loop A (5’ SLA) in the 5’UTR to initiate the synthesis of negative strand RNA (Filomatori et al. 2006). Recent structural and biochemical studies of dengue SLA binding with NS5 implicated K22/S23 and K841/R842 responsible for the binding (Lee et al. 2022). While K22/S23 are not well conserved among flavivirus and could tolerate diverse mutations, K841/R842 shows strong amino-acid preference toward amino acids with positive charges (Figure 4A). This result suggests that K841/R842 interaction with 5’ SLA could be crucial to DENV replication. A previous study has implicated R770, R773, Y838, K841, and R856 as the residues responsible for the interactions with stem loop at the 3’end (3’ SL) (Hodge et al. 2016). These residues are conserved across flavivirus and have strong amino-acid preferences for positive-charge amino acids or aromatic amino acid (Y and F for residue 838) (Figure 4A).

**Figure 4:**
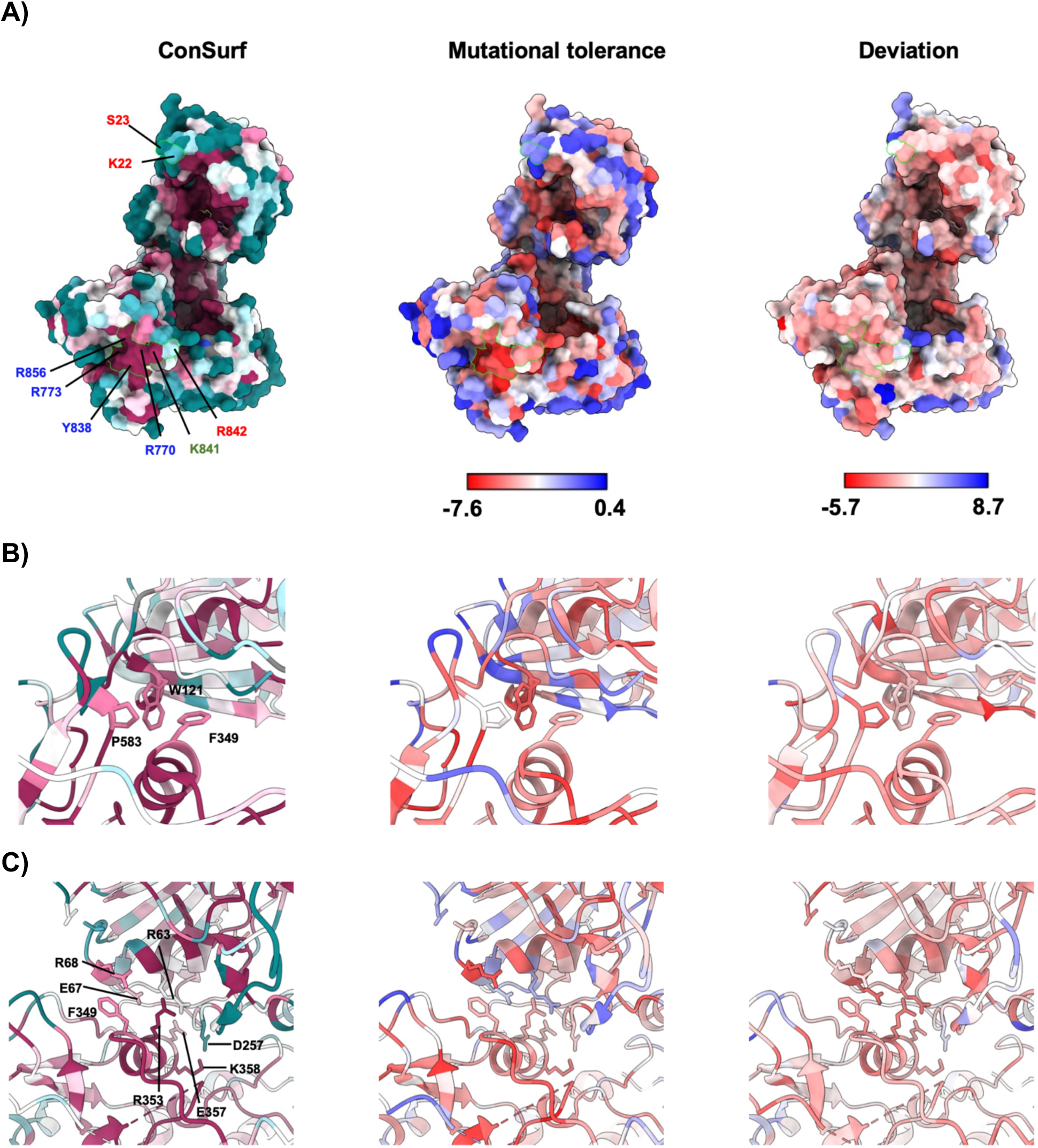
Structural mapping of ConSurf score, mutational tolerance, and sequence conservation of functional sites on RdRp domain. A) Structural mapping on 6kr2.pdb with highlighted residues that interact with 5’ SLA (labeled in red) and 3’ SL stem loops (labeled in blue) with K841 (labeled in green) shared by both noncoding RNA elements. B) Structural mapping of interacting residues between the MTase and RdRp in JEV conformation (6kr2.pdb, Wu et al. 2020). C) Structural mapping of interacting residues between the MTase and RdRp in DENV3 conformation (6kr3.pdb, Wu et al. 2020).

Previous studies have shown that the interactions between MTase and RdRp domains enable efficient viral RNA synthesis by flavivirus RdRp (Wu et al. 2015, Zheng et al. 2022, Li et al. 2014, Wang et al. 2012,). Several crystal structures of dengue NS5 have captured two distinct conformations of the interdomain interactions, defining the residues involved (Lu et al. 2013, Zhao et al. 2015b, Sahili et al. 2019, Wu et al. 2020). In the extended JEV conformational mode, W121 from MTase and F349, P583 from RdRp form hydrophobic interactions in DENV2 NS5 (Wu et al. 2020). W121 shows strong amino-acid preference and does not tolerate mutations (Figures 2A and 4B). F349 show strong amino-acid preference for aromatic residues. (Figures 2A and 4B). However, P583 could tolerate several mutations (Figures 2A and 4B). In the DENV3 conformational mode, E67, R68, R63, F349, R353, E357, K358 forms the interdomain interactions (Wu et al. 2020). R68 has strict preference for R, while F349 prefers aromatic F,Y,W and leucine. R353, E357, K358 strongly prefer the amino acids that maintain their respective charges. However, R63 and E67 have no clear amino-acid preferences (Figures 2A and 4B). As the majority of the interacting residues have strong amino-acid preferences, our data agree with the current model that both conformational modes are important for flavivirus replication.

### Analysis of catalytic site of NS5 MTase domain

Flavivirus MTase catalyzes three steps of viral RNA capping and methylation. First, its guanylyltransferase activity (GTase) transfer GMP moiety from GTP to the ppN-RNA to generate GpppN-RNA cap structure. Second, its N7 MTase activity transfers methyl group to N7 position of the G cap to create m^7^GpppN-RNA. Lastly, its 2’-O-MTase activity methylates the 2’-O position of the first nucleotide after the m^7^G cap to generate m^7^GpppNm-RNA (cap-1 structure). SAM serves as the methyl donor for both N7 and 2’-O methylations. N7-methylation is required for flavivirus viability as it is needed for translation initiation. 2’-O-methylation is not required but is needed for efficient virus replication in type I-IFN competent cells (Zhou et al. 2007, Daffis et al. 2011).

Several structures of flavivirus MTase have shown that F25 is the key residue that anchor the G cap throughout the three steps of the viral RNA capping and methylation. While this residue is strictly conserved among the flavivirus, our DMS showed that it could be exchanged with Y or W which also have side chains with aromatic rings, indicating that *π*-*π*stacking is required for fixing the G cap rather than the F per se (Figure 5A). R/K 29 is the key residue shown to form the GMP-arginine adduct, an intermediate during G transfer from GTP to the ppN-RNA. While previous biochemical and structural studies showed that R/K29 covalently binds to GMP (Bollati et al. 2009, Issur et al. 2009, Jia et al. 2022), our DMS showed that A, S, and C could also be accommodated. Nevertheless, the fitness of R29A, R29S, and R29C were lower than that of R29K (Figures 2A, 5A). As the side chain of R29A would not provide functional group capable of forming a covalent bond with GMP, the viability of R29A raises the question whether flavivirus MTase could perform guanylyl transfer through an alternative, yet less efficient pathway. Similarly, P152 and S214 that were shown to interact with GpppN cap structure could be mutated to other amino acids despite their high conservation among flavivirus (Figures 2A, 5A). Among the five key cap-interacting residues, R212 was the only exception with both MSA and DMS showing strict conservation and absolute amino-acid preference, suggesting that the interaction between its guanidinium group and *β*-/*γ*-phosphates of the GTP substrate could be essential for the catalysis of guanylyl transfer (Figure 5A).

**Figure 5:**
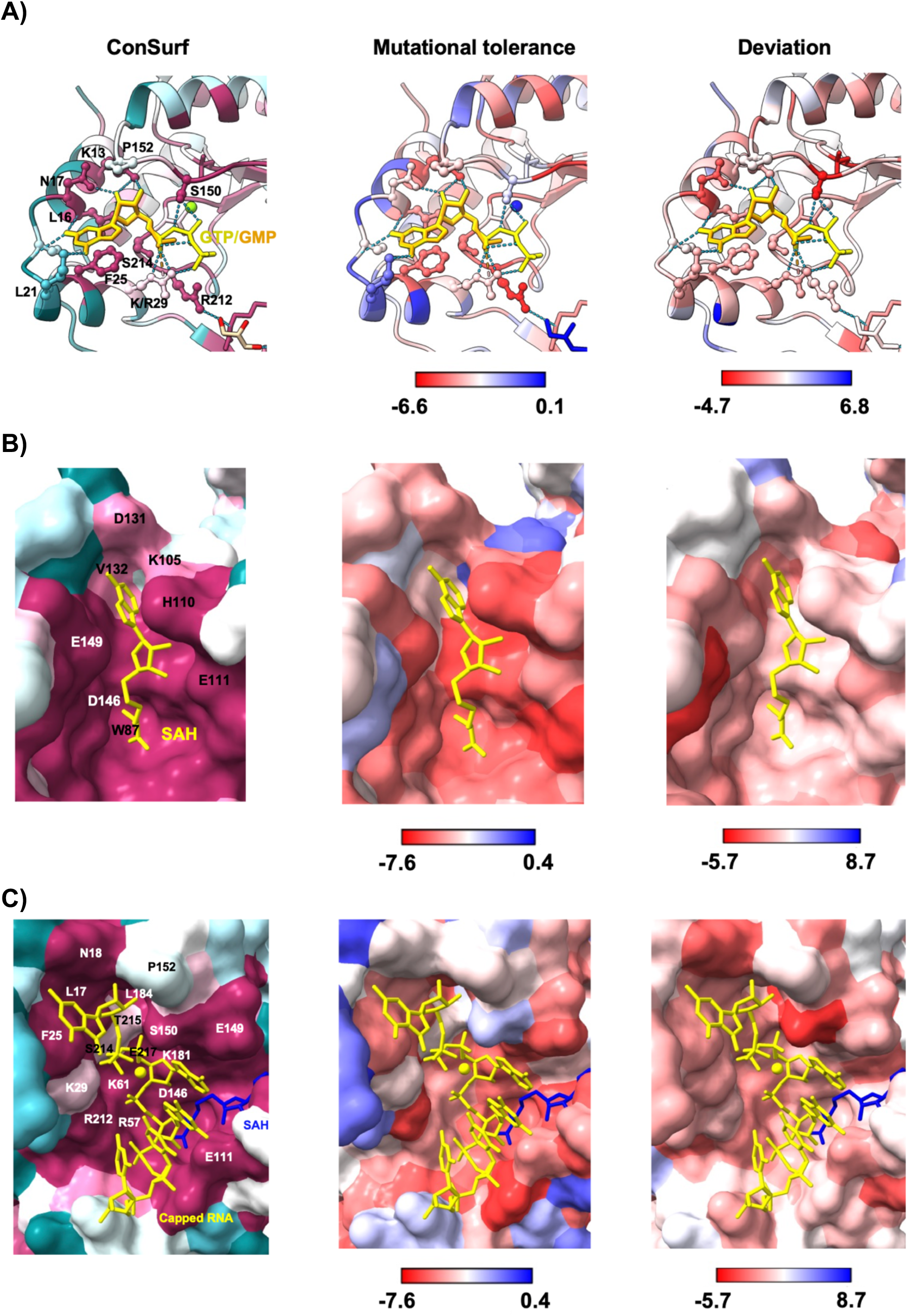
Analysis of mutational tolerance and sequence conservation of the MTase active site. A) Structural mapping of ConSurf score, mutational tolerance, and deviation score on the GTase active site using Omsk hemorrhagic fever virus (OHFV) MTase structure (7v1g.pdb, Jia et al. 2022). B) Structural mapping on the SAM binding site on 6kr2.pdb (Wu et al. 2020). C) Structural mapping on the ternary complex of DENV3 MTase + capped RNA (m^7^GpppAGUU) + SAH (5dto.pdb, Zhao et al. 2015a). Residues are numbered by DENV2 NS5 positions.

The residues that make up SAM pocket has relatively low mutation tolerance (Figure 5B). H110, a key residue that form hydrogen bonding with adenosine, showing strict conservation and absolute amino-acid preference. D131 could accommodate E and N which have the side chains that could still maintain hydrogen bonding with the adenine base. Similarly, W87 could accommodate F and Y with aromatic side chains. V132 could accommodate several hydrophobic amino acids despite its strict sequence conservation. D146, G86, and S56 has strong amino-acid preference and strict sequence conservation. Previous biochemical study showed decoupling of N7- and 2’- O-methylation through mutations in the SAM pocket, with lethal mutations affecting N7- methylation (Kroschewki et al. 2008). The viability of the mutants targeting these residues are consistent with our DMS data (Figure 2A).

Flavivirus MTase relies on a K-D-K-E tetrad (K61-D146-K181-E217 for DENV2) for both N7 and 2’-O methylation. Previous studies showed that mutations to the each of the K-D-K-E residues had differential effects on the N7 and 2’-O-methylations (Zhou et al. 2007). K61, D146, K181, and E217 show strong amino-acid preferences toward the wild-type amino acids (Figure 2A). As our DMS assayed basal mutant fitness in mammalian cells compromised in type I IFN signalling, the mutational effects on 2’-O-methylation may not be reflected in the data. Nevertheless, most residues that interact with viral RNA in a 2’-O-methylation conformation have relatively low mutation tolerance, consistent with active site sharing between N7 and 2’-O-methylation (Figure 5C) (Dong et al. 2010a and 2010b). Notably, S150 is tolerant to mutations despite its strict sequence conservation (Figures 5A, 5C).

## Discussion

Our DMS analysis of dengue NS5 in DENV2-16681 genetic background has provided mutational database useful for understanding the functions of flavivirus NS5. The effect of saturation mutagenesis at each residue could be used in combination with structural and biochemical data to examine the chemical natures of key residues involved in its enzymatic functions, revealing deeper insights than conventional alanine scanning used to study flavivirus NS5. Our DMS data suggests that priming loop could have more roles than priming viral RNA synthesis. The unexpected mutation tolerance of multiple residues in the highly conserved RdRp active site suggests a possibility of engineering the active-site motifs to modulate error rate of viral RNA synthesis. The mutational tolerance at K29 in the MTase domain suggests that flavivirus NS5 might be able to utilize an alternative pathway to the R/K29-GMP adduct formation to transfer guanylyl group to the 5’ end of viral RNA.

While we have focused the analysis of our DMS data on the enzymatic functions of NS5 here, the DMS dataset could also provide a general framework for genetic engineering of flavivirus for live-attenuated vaccine development and future investigations of other functional aspects of flavivirus NS5. It could complement structural information in rational design of new antiviral drugs to minimize risk of viruses developing drug resistance. The DMS dataset could be useful in modeling the molecular dynamics of NS5 to identify potential conformational changes relevant for its essential functions. It will be useful for interpreting the structures of NS5 functions at different steps of viral RNA synthesis as more structural and biochemical data of those steps become available.

The DENV2-16681 and the NS5 gene libraries could be analyzed in different settings to dissect NS5 functions in immune response and evolution. The libraries could be probed in mosquito cells to dissect flavivirus-mosquito interactions, potentially providing insights into flavivirus evolution to infect diverse insect vectors. Analyses of infection in the presence of type-I IFN and NS5 interactions with key regulators such as STAT2 and PAF1C with the libraries could offer ways to dissect NS5 and type-I IFN signaling, potentially revealing novel attenuation mutations that specifically target virus immune-antagonistic functions (Wang et al. 2020a, Petit et al. 2021). DMS analysis of influenza viruses with type-I IFN have identified novel mutations that could enhance flu vaccine efficiency (Du et al. 2018). The libraries could be used to probe NS5 interactions with antiviral small molecules that target allosteric sites (e.g. the N pocket (Lim et al. 2016, Gharbi-Ayachi et al. 2020), A and B cavivities (Zou et al. 2011), RNA tunnel (Niyomrattankit et al. 2010, Arora et al. 2020) of NS5 to dissect its conformational dynamics essential for its functions.

## Material and methods

### Cell lines

BHK21-rtTA3 cell line was maintained in DMEM supplemented with 10% heat-inactivated fetal bovine serum (FBS), penicillin/streptomycin, sodium pyruvate, and high glucose (HyClone). Vero cells were maintained in DMEM supplemented with 10% FBS and penicillin/streptomycin. A549 cells were maintained in R10 (RPMI 1640 supplemented with 10% FBS, penicillin/streptomycin, and 1x MEM non-essential amino acids). Cells were cultured at 37°C, 5% CO_2_ concentration, and 95% relative humidity.

### Infectious titer quantitation

Infectious titers of DENV were quantitated by foci assay with 4G2 monoclonal antibody according to the protocol detailed in Siridechadilok et al. (2013) (Siridechadilok et al., 2013)

### Construction of NS5 gene library

NS5 gene library was first created with the high-copy plasmid pUC57 for the construction of DENV2 library (summarized in Supplementary Figure 1A). DENV2-16681 NS5 gene was first cloned onto pUC57. To design synthetic oligonucleotide library for NS5 library construction, NS5 was broken down into 45 frames for mutagenesis, with each frame covering 20 residues. In addition to the target 20 residues for mutagenesis, each frame also included unmutagenized flanking sequences for DNA assembly during gene library construction. For each amino-acid mutation, the codon with highest codon usage in human was used for designing mutagenic oligonucleotide to minimize the size of the library. Focusing on fixed, defined codons instead of randomized NNN or NNK codons should also reduce sequencing error propagation during read-count analysis as the error from one codon would not always end up counted in the other codons. 1-2 synonymous mutations at each residue (with the exception of 1-codon methionine and tryptophan) were also incorporated into the library as controls for the screen. A total of 18,433 genetic variants were designed in the oligo pools used for NS5 library construction. The mutagenic oligonucleotide library was splitted into two sets of odd and even frame numbers so that no adjacent frames were included in each set to prevent concatenation of frames during PCR amplification. The two sets of oligo libraries were synthesized on two separate chips by GenScript. The mutant sequences of each frame were amplified with Phusion DNA polymerase (Thermo Scientific) by a pair of primers that bound to the flanking sequences on the oligonucleotides. The PCR products for each frame was gel-purified and assembled with the PCR products of the pUC57-NS5 that was amplified with the reverse pair of flanking primers (containing the sequence of the whole plasmid with the target mutagenic region left out). DNA assembly was done with Gibson Assembly (Gibson et al. 2008). The assembly was electroporated into 10B *E.coli* (NEB). The library construction of pUC57-NS5 was performed for each frame separately, creating a total of 45 pUC57-NS5 plasmid libraries (Supplementary Figure 1A).

### Construction of DENV2 infectious-clone library

The infectious-clone plasmid construct of DENV2 with an intron insert will be detailed elsewhere (Suphatrakul et al., manuscript in preparation). To generate DENV2 infectious clone plasmid library, the NS5 gene from pUC57-NS5 plasmid library was amplified by PCR. The PCR product was then assembled with the backbone generated from the DENV2-16681 infectious-clone plasmid by a PCR amplification that left out the NS5 gene (Supplementary Figure 1B). The backbone and the NS5 PCR library were assembled by Gibson assembly (Gibson et al. 2008). The assembly reaction was cleaned up by AMPure XP magnetic bead (Beckman Coulter) and electroporated into DH5a. The electroporated DH5a was cultured at 30°C in LB-Amp broth. The bacteria were harvested for plasmid library preparation. DENV2 infectious-clone plasmid library of each NS5 frame was constructed individually. Two consecutive library frames were combined into a single pool to make 23 pools (except for the 23^rd^ pool that contains only the 45^th^ frame) of the infectious-clone plasmid library.

### Deep mutational scanning

Each pool of infectious-clone plasmid library was transfected into BHK21-rtTA3. BHK21-rtTA3 was treated with 1μg/ml doxycycline during seeding and maintained after transfection. (Suphatrakul et al. 2018). The media was harvested 3 days post transfection and transferred to infect confluent Vero cells. The final DENV2 pool libraries were harvested twice at 3 and 5 days post infection. Each DENV2 pool library was precipitated with 10% PEG8000 overnight at 4°C and harvested by centrifugation. The pellet was resuspended in 1xPBS and supplemented with 20% FBS for storage. Three replicates of DENV2 libraries were generated and deep sequenced. One million infectious units (FFUs) of virus pool stock was used for viral RNA extraction by GeneAid viral RNA extraction kit (GeneAid) and converted to cDNA using ImProm-II™ reverse transcription system (Promega) according to manufacturers’ protocols. The target NS5 region with mutations was amplified by PCR with Phusion DNA polymerase (Thermo Scientific). The gel-purified PCR products of all pools were combined together into one sample for deep sequencing. Deep sequencing was performed by Novaseq with 150-bp paired-end read length.

### DMS data analysis

We used DiMSum wrapper to pre-process the fastq data using default settings (Faure et al. 2020). We generated the final count of each mutant using a custom python script. We performed read count analysis in two ways. First, we counted mutants by including all the possible NNN codons. Second, we counted mutants by including the sequencing reads that exactly matched the designed oligo libraries. Classification of the input read counts by the number of nucleotide mismatches to the wild-type sequencing reads showed a distinct peak that differentiated the histograms of the NNN-codon and the fixed-codon read counts (Supplementary Figure 2C). Using the classified histograms, we set the input read-count thresholds to 60, 25, and 25 for mutants with the Hamming distances of 1, 2, and 3 nucleotides, respectively. From the read counts, we re-created the vsearch.unique files of the input and the outputs with full-length NS5 sequences for fitness score calculation with normalization and error model fitting by STEAM analysis in DiMSum (Supplementary Figure 3). Amino-acid preference was calculated using dms_tools2 with `--method ratio` option (Bloom, 2015). Mutation tolerance of each residue was calculated by averaging the weighted-average fitness scores of the nonsynonymous mutants that passed the read-count filter.

### Validation of the fitness of DENV2 mutants

DENV2 mutant viruses were constructed based on the protocol in Siridechadilok et al. 2013. The pGEM-DENV2 infectious clone plasmid was used as the template for backbone PCR. The NS5 mutant genes were created by assembling two pieces of PCR products where the mutations were introduced by a mutagenic primer at the assembly joint. The complete infectious-clone plasmid assembly, thus, was assembled from three PCR products. The mutant viruses were generated by transfection of the assembly into BHK21-rtTA3 and expanded in Vero cells to generate stocks. The virus mutations were confirmed by Sanger sequencing of the NS5 gene. The infection in A549 was carried out at MOI = 0.1 for 2 hours at 37°C and washed twice afterward by 1xPBS. The infected cells were cultured in R10 for 48 hours before harvest for titration.

### Multi-sequence alignment of flavivirus NS5

Multi-sequence alignment (MSA) of NS5 was done with MAFFT version 7 (Katoh and Standley, 2013) using 95 non-segmented flavivirus representative NS5 sequences commissioned by the International Committee on Taxonomy of Viruses (ICTV, Supplementary Figure 4A). The MSA was displayed using MView (https://www.ebi.ac.uk/Tools/msa/mview/, Brown et al. 1998) and WebLogo plot (https://weblogo.berkeley.edu/logo.cgi, Crooks et al. 2004) (Supplementary Figure 4C). Phylogenetic tree based on the NS5 sequences (Supplementary Figure 4B) were constructed using the following setting on MAFFT web server: neighbor-joining algorithm, conserved sites (758 AAs), substitution model = JTT, ignore heterogeneity among sites, and bootstrap on with 100 number of resampling. ConSurf scores were calculated using the MAFFT alignment file on ConSurf webserver (https://consurf.tau.ac.il/, Ashkenazy et al. 2016). For displaying on NS5 structures, the ConSurf scores were partitioned into 9 grades as detailed on the ConSurf web site (bin 9 contains the most conserved positions and bin 1 contains the most variable positions). Deviation score for each residue was calculated as the difference between the mutation tolerance predicted by the linear equation derived from fitting the mutation tolerance and the ConSurf scores (Figure 2C) and the mutation tolerance observed from the DMS data.

### Structural analysis

Projection of ConSurf scores, mutational tolerance, and deviation onto crystal structures were done with PyMol and displayed using ChimeraX version 1.3 (Pettersen et al. 2021).

## Data availability

The PDB files with mapped mutational tolerance and deviation scores are provided as supplementary files. The csv table of the averaged fitness and the standard error of each single amino-acid mutant have been deposited in MaveDB database (Esposito et al. 2019) under accession number <u>urn:mavedb:00000111-a-1</u>. The raw deep sequencing data have been deposited in the Sequence Read Archive under BioProject accession number PRJNA941815.

## Acknowledgement

B.S. would like to acknowledge the support from Newton-NSTDA- MRC program in infectious diseases and the Grand Challenges Canada. J.M. would like to acknowledge the support from MRC, UK. B.S., A.S., P.P., N.S., R.N., and S.O. have filed a patent application related to this work. We would like to thank Andre Faure for his help with DiMSum; Pala Chidpratum for technical assistance.

---

**Supplementary Figure 1:**
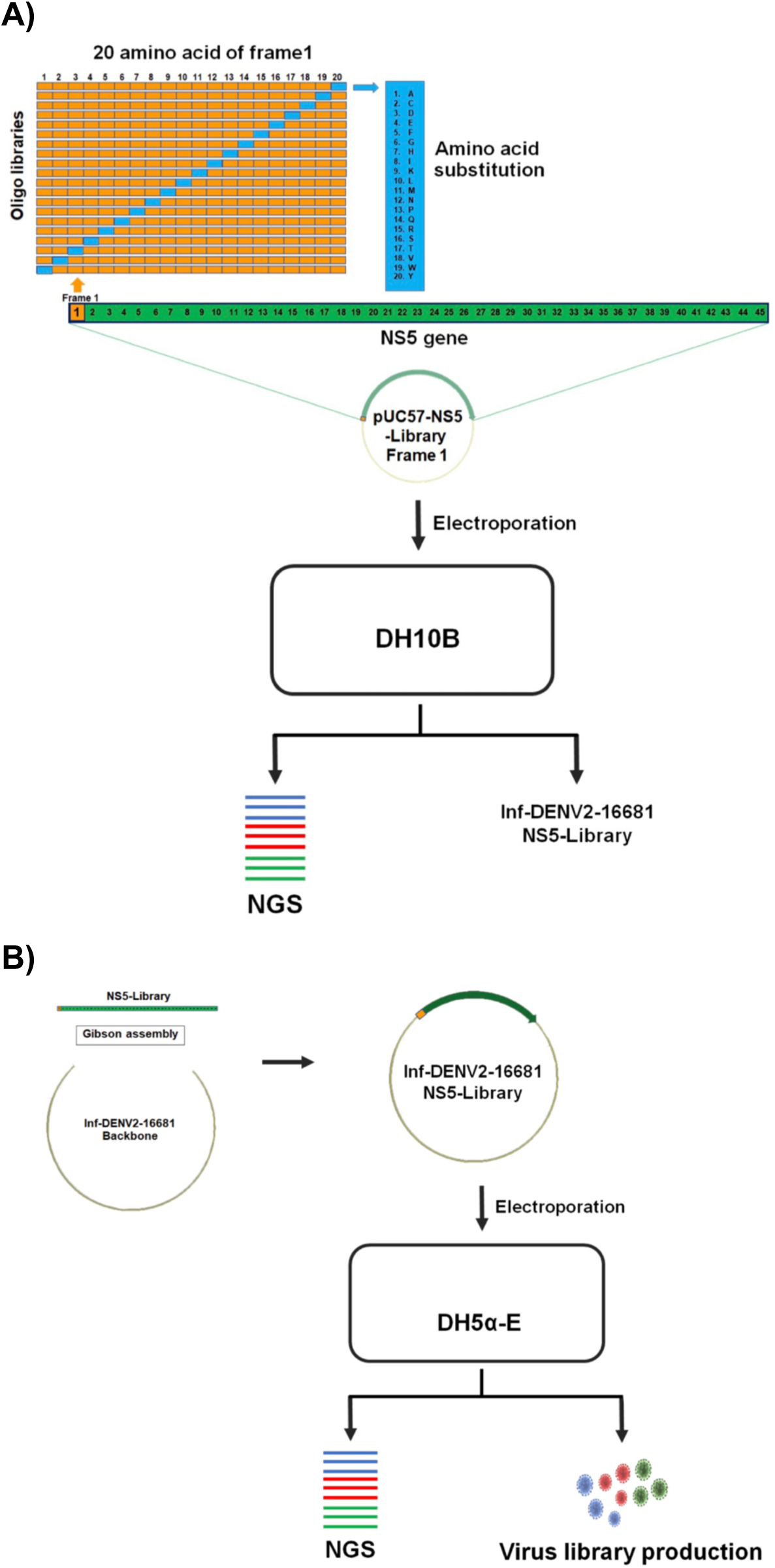
Construction of pUC57-NS5 plasmid library and DENV2- NS5 infectious-clone plasmid library. A) Construction of pUC57-NS5 library from oligonucleotide libraries. B) Construction of infectious-clone plasmid library from the pUC57-NS5 library.

**Supplementary Figure 2:**
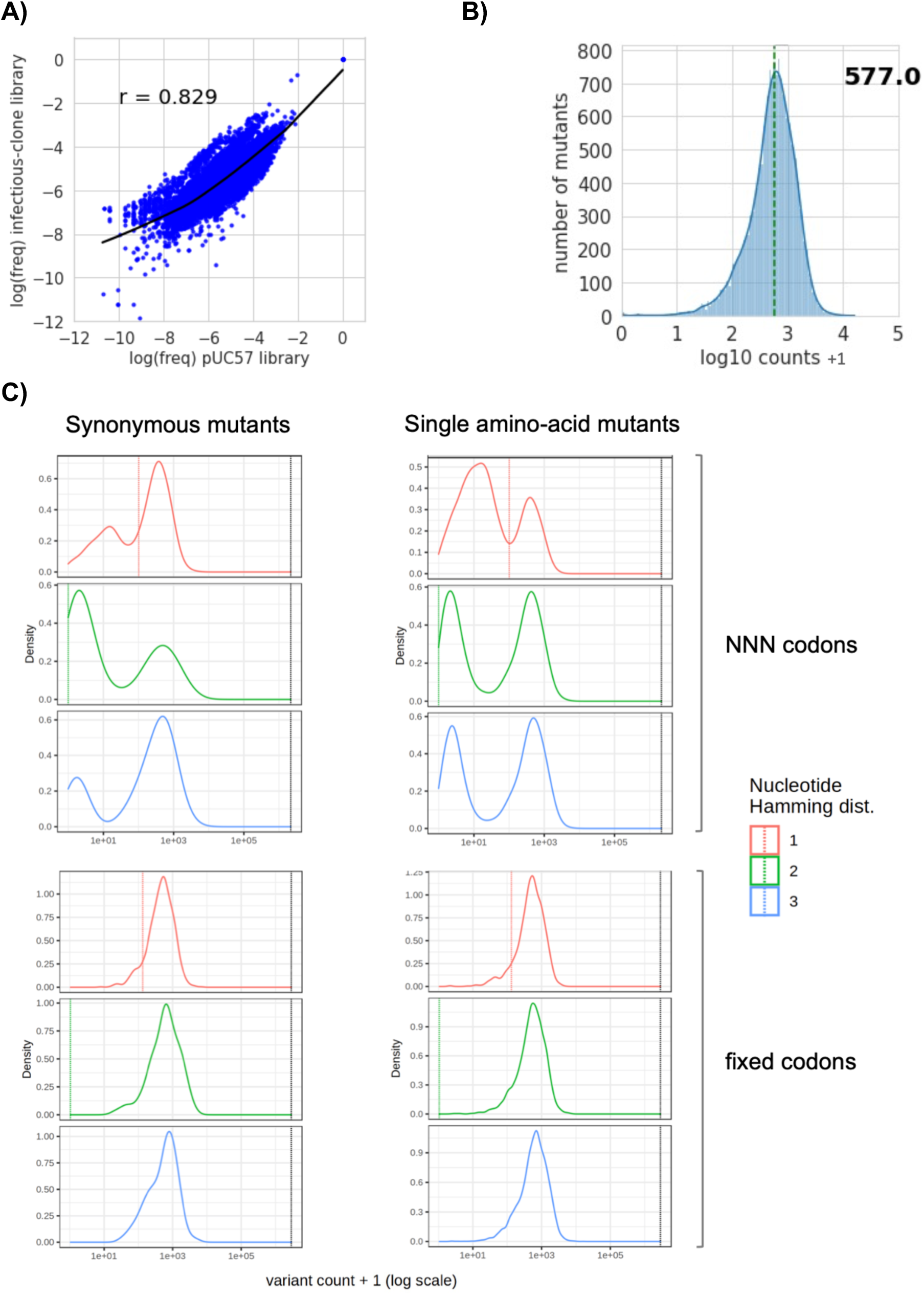
Statistical analysis of pUC57-NS5 plasmid library and DENV2-NS5 infectious-clone plasmid library. A) The correlation between the log (mutant frequency) of designed mutants (both synonymous and nonsynonymous) in pUC57 and infectious-clone plasmid libraries. B) Histogram of read counts of NS5 mutants in the infectious-clone plasmid library. The number shown is the average read counts. C) Histograms of read counts of designed (or fixed codon) mutants (synonymous and nonsynonymous) and the NNN mutants of the infectious-clone plasmid library, categorized by 1, 2, and 3 nucleotide differences.

**Supplementary Figure 3:**
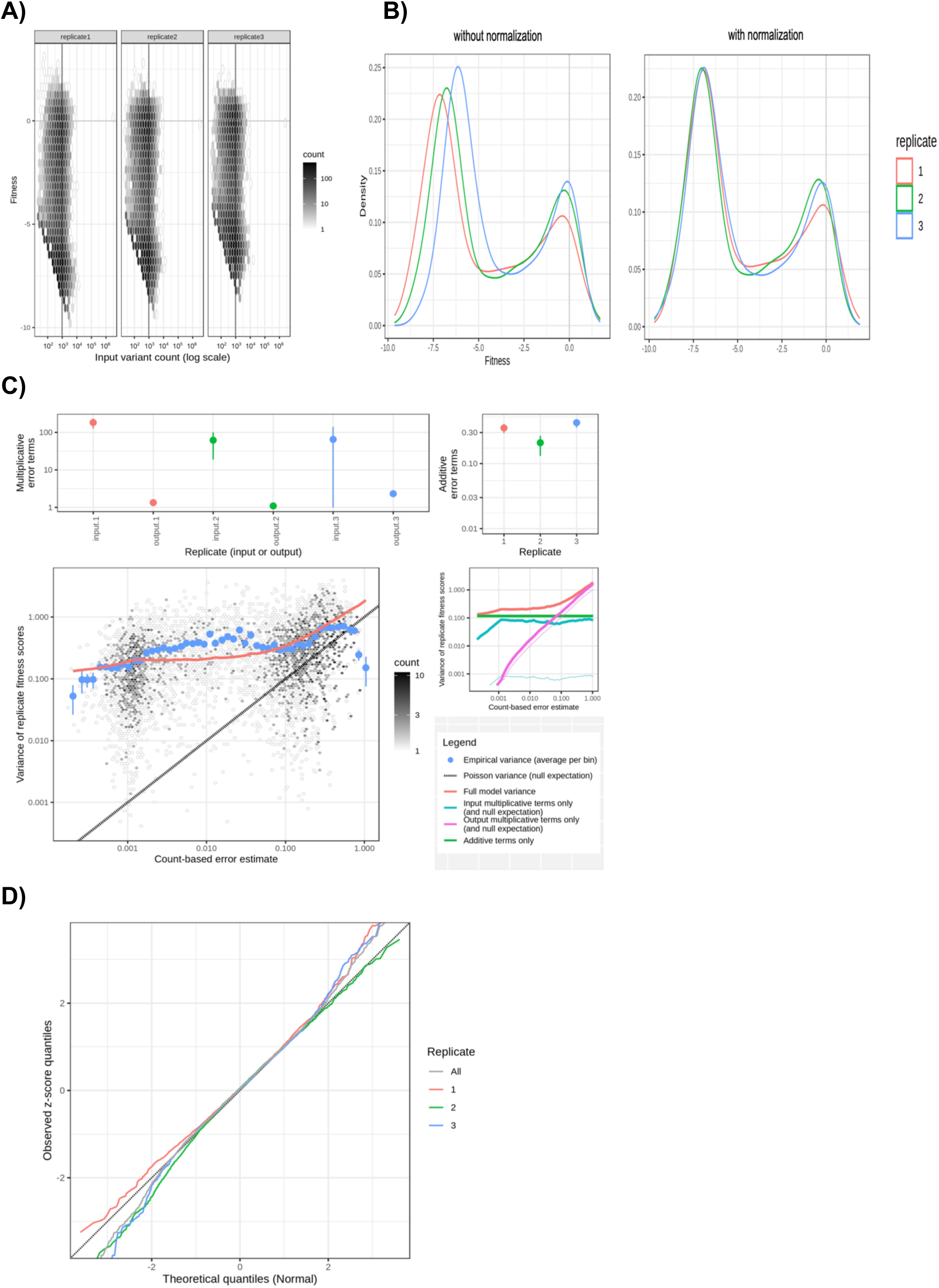
Statistical analyses and error estimates by DiMSum. A) Distribution of mutant fitness and input read counts. Input count threshold above which variants are expected to span the full fitness range (y-axis spread) is displayed as black vertical dashed line. Variants surpassing this threshold are used to fit the error model. B) Fitness distribution of each replicate before and after normalization. C) Fitness error model fitting by DiMSum. The upper panels show multiplicative (upper left panel) and additive (upper right panel) error terms estimated by the DiMSum error model. Dots give mean and error bars indicate the standard deviation of parameters over 100 bootstraps. The lower left plot shows variance of fitness scores between replicates as a function of sequencing count-based (Poisson) variance expectation (average across replicates). The black dashed line indicates perfect correspondence (i.e. Y=X). The full DiMSum error model (red line) describes deviations from the null expectation (black dashed line) in the observed variance of fitness scores. The lower right plot compares the full DiMSum error model (red line) to variance contributions using either input multiplicative error terms (cyan line), output multiplicative error terms (magenta line) or additive error terms (green line) only. Dashed cyan and magenta lines indicate purely sequencing count-based variance expectation corresponding to input and output samples, respectively. D) The quantile-quantile (Q-Q) plot assessing the performance of the fitness error model using leave-one-out cross-validation.

**Supplementary Figure 4:**
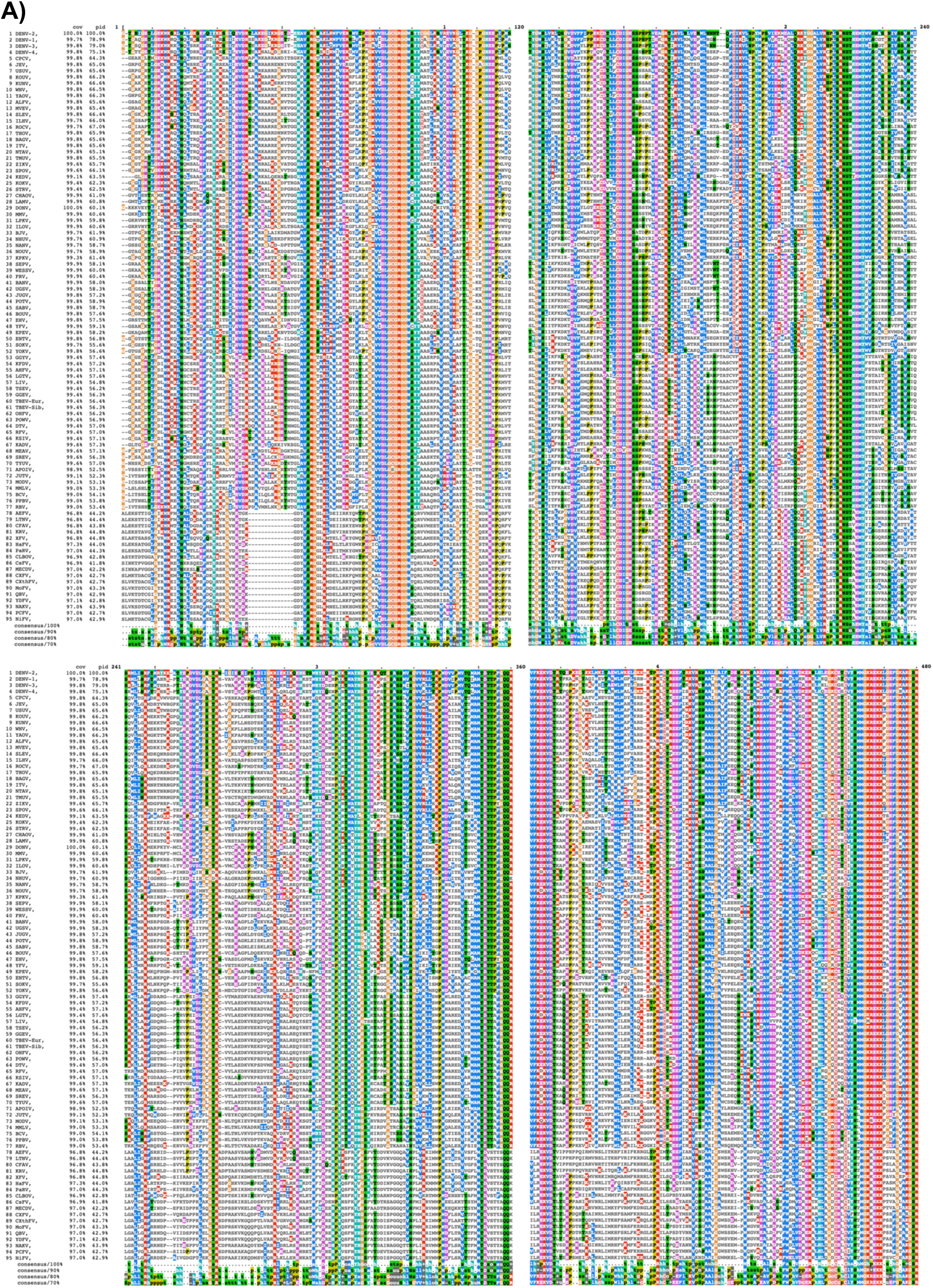
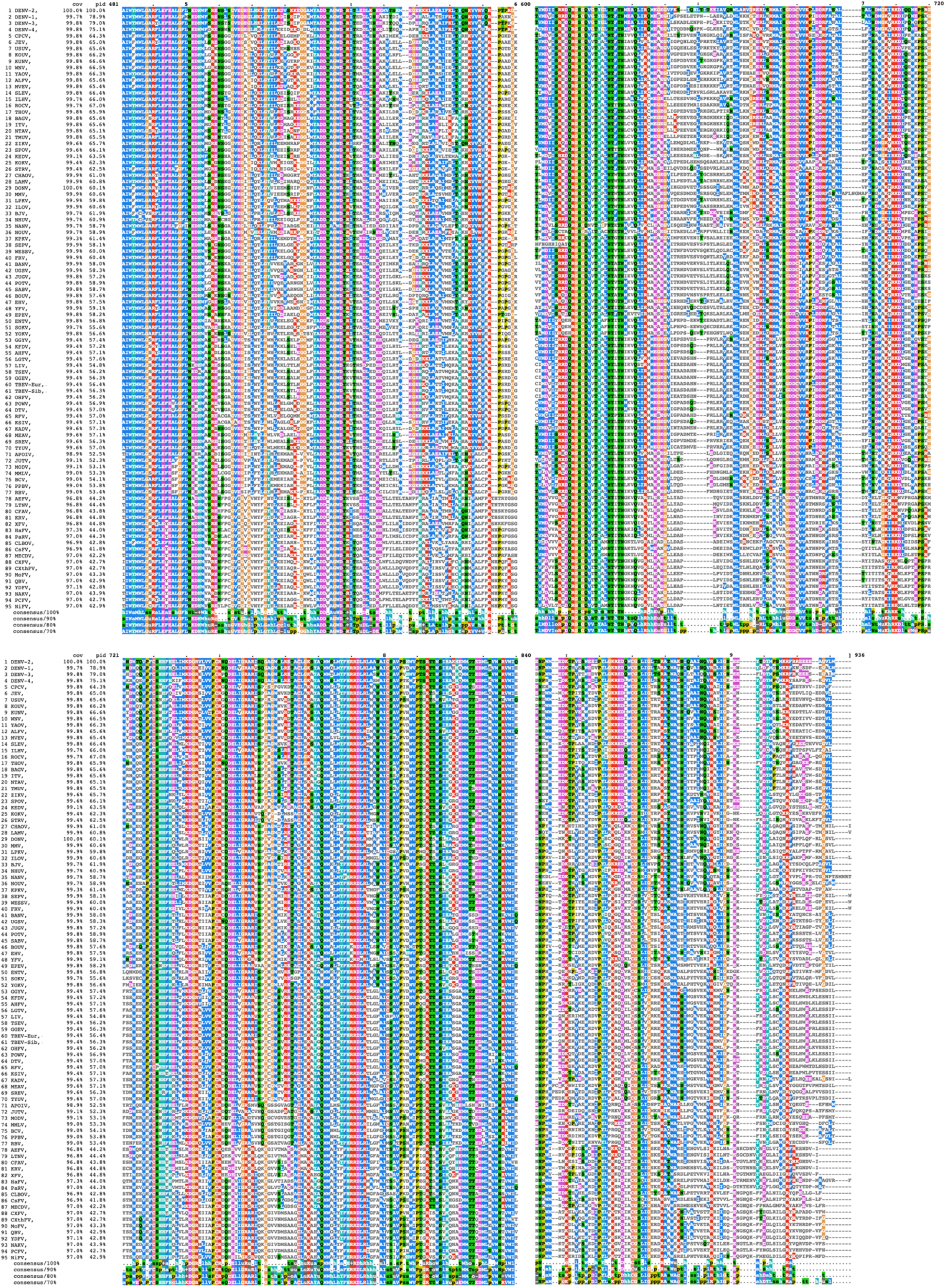
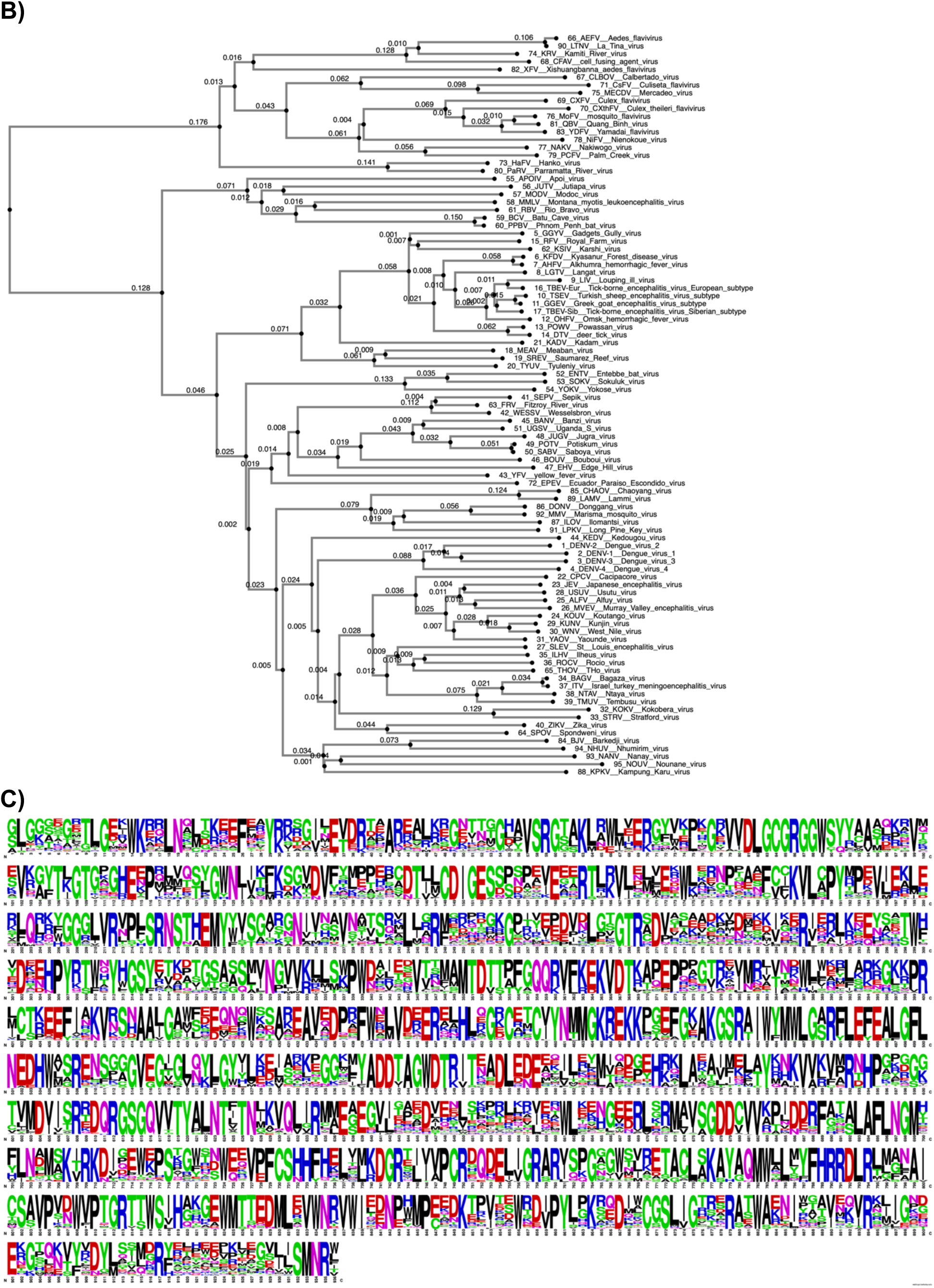
Multiple sequence alignment (MSA) of non-segmented flavivirus NS5 by MAFFT. A) MSA of 95 representative flavivirus NS5. B) Phylogenetic tree based on the MSA of the representative flavivirus NS5. C) WebLogo plot of NS5 MSA.

